# Effect of endogenous earthworm’s extracts on “plant growth-promoting fungi”

**DOI:** 10.1101/2024.11.14.623545

**Authors:** Lamia Yakkou, Sofia Houida, Mohammed Raouane, Musa Hatipoğlu, Souad Amghar, Abdellatif El Harti, Mohammed Jaber

## Abstract

This study explores the in vitro effects of earthworm-derived crude extracts on the growth of key Plant Growth-Promoting Fungi (PGPF) strains, including *Trichoderma* sp., *Melanocarpus* sp., and *Acaulospora* sp. Crude extracts were prepared from freshly harvested earthworms (HE) and those that had been starved and devoid of soil for 10 days (FE), and their effects on fungal proliferation were assessed at varying concentrations. Results demonstrated that both HE and FE extracts significantly enhanced fungal growth rates compared to traditional growth media (PDA and SAB), even at lower concentrations. The enriched nutrient profile and growth factors present in these extracts suggest that earthworm decomposition contributes valuable bioactive compounds to the soil, promoting PGPF activity and soil fertility. Additionally, media derived from these extracts stimulated rapid spore germination, which is critical for fungal dissemination and ecological establishment. Notably, the study found that contents within the earthworm digestive tract did not influence fungal growth, affirming that the growth-promoting effects are inherent to the earthworm tissue structure itself. These findings underscore the potential of earthworm-based media as a natural and effective growth medium for enhancing PGPF propagation, with implications for sustainable agricultural practices and soil health improvement.

## Introduction

Soil is regarded as a dynamic, living system, shaped by the intricate interactions among microorganisms and fauna that regulate the availability of plant nutrients. As noted by Prasad et al. (2019), agricultural productivity cannot be sustained without the presence of a diverse microbial community within the soil. Among the most promising microbial agents for enhancing soil health are Plant Growth-Promoting Fungi (PGPF) (Zainab et al., 2021). Extensive research has demonstrated that PGPF serve as a viable alternative to chemical fertilizers, offering a more environmentally friendly and sustainable approach to agriculture. Rhizosphere soils enriched with various PGPF taxa have been shown to promote plant growth both directly and indirectly (Thakor et al., 2016). These fungi contribute to plant development by mobilizing soil nutrients, producing various plant growth regulators, providing protection against pathogens, and facilitating the breakdown of xenobiotics (Zin and Badaluddin, 2020). Additionally, numerous studies have highlighted the efficacy of PGPF species, such as those from the *Ampelomyces, Coniothyrium*, and *Trichoderma* genera, in mitigating both biotic and abiotic stressors (Beneduzi et al., 2012; Rozier, 2017; Zin and Badaluddin, 2020).

Recent studies have highlighted the beneficial effects of interactions between earthworms and PGPF (Zarea et al., 2009). For example, the inoculation of *Pheretima sp*. earthworms with the fungus *Glomus mosseae* resulted in improved forage yield, enhanced mycorrhizal colonization, increased nitrogenase activity in free-living rhizosphere bacteria, elevated soil microbial biomass carbon, and enhanced growth of clover plants (Zarea et al., 2009). Earthworms are recognized as “ecosystem engineers” due to their significant impact on soil properties and the structure and activity of microbial communities (Zhao et al., 2010; Bernard et al., 2012; Al-Maliki et al., 2020). These annelids have shown selective preferences for certain fungal species in the soil (Moody et al., 1996; Gómez-Brandón and Domínguez, 2014). However, the relationship between earthworms and fungi is complex. Previous research has indicated that earthworms tend to prefer rapidly growing fungal species, particularly those associated with the early stages of decomposition, such as cellulolytic fungi and those consuming soluble carbohydrates (e.g., *Trichoderma* spp.) (Moody et al., 1996; Bonkowski and Schaefer, 1997). Anecic earthworms, in particular, have been shown to favor litter colonized by specific fungal species, including *Fusarium lateritium* and *Trichoderma* spp. (Moody et al., 1996). The survival of fungal spores within earthworms, however, is highly dependent on the characteristics of the spores. For instance, *Fusarium lateritium* spores did not survive passage through the intestines of *Lumbricus terrestris* and *Aporrectodea longa* (Brown, 1995; Moody et al., 1996). In contrast, spores of *Chaetomium globosum* passed through the intestines of both earthworm species without affecting their viability. Furthermore, certain fungal species, particularly toxin-producing genera such as *Aspergillus, Fusarium*, and *Penicillium*, can be detrimental to earthworms (Edwards and Fletcher, 1988; Morgan and Morgan, 1988). Bonkowski et al. (2000) conducted preference experiments to assess earthworm species’ selection for various soil fungi. They concluded that earthworms use early-successional fungal species as indicators of fresh organic matter, although the underlying mechanisms driving this preference remain unclear.

As they navigate through soil, earthworms deposit cutaneous mucus and, over time, leave their bodies, both of which are metabolized by soil microorganisms and contribute to plant-available nutrients. These saprophagous organisms represent a substantial biomass, ranging from 0.5 to 5 tonnes per hectare (fresh weight) (Edwards and Fletcher, 1988; Lavelle, 1988; Hameed et al., 1994). Their remains can significantly enrich soil with endogenous nutrients and stimulate microbial activity. Previous studies have reported high crude protein content in various earthworm species: *Lumbricus rubellus* contains 65.63% (Damayanti et al., 2008), *Lumbricus terrestris* 32.60% (Julendra et al., 2010), and *Perionyx excavatus* 57.2% (Tram et al., 2005). In *Eisenia foetida*, proteins comprise 61.85%, fats 11.13%, and ash 8.7% of the dry matter (Medina et al., 2003). Additionally, mucus from anecic species (*Lumbricus terrestris*) and endogeic species (*Aporrectodea caliginosa*) is predominantly composed of proteins and carbohydrates (Zhang et al., 2010).

Our laboratory has focused on examining the in vitro post-mortem effects of earthworm crude extracts on the growth of soil-beneficial PGPF. We analyzed fungal development after exposing strains to various concentrations of earthworm crude extracts. The fungal species utilized (*Trichoderma* sp., *Melanocarpus* sp., *Acaulospora* sp.) are recognized as PGPF, given their crucial roles in enhancing soil fertility and promoting plant growth, both directly and indirectly (Clark, 1997; Naseby et al., 2000; Siddiqui and Shaukat, 2004; Zin and Badaluddin, 2020).

## Materials and Methods

### Earthworms and Fungi

The earthworm species *Aporrectodea molleri* was collected from the Akrach region, located approximately 20 km southeast of Rabat, Morocco. This species is indigenous to the area and dominates the local earthworm population. Three fungal species were tested: *Trichoderma sp*., *Melanocarpus sp*., and *Acaulospora sp*., all of which are part of the laboratory collection of the Earthworms, Soil Productivity Improvement, and Environmental Studies Laboratory (LAPSE) in Rabat, Morocco. These strains were identified through amplification and sequencing of the ITS gene, with confirmation provided by the National Center for Scientific and Technical Research (CNRST) in Rabat. All three fungi belong to genera known for their beneficial roles in soil health.

### Preparation of culture media from raw extracts of earthworms

#### Preparation of raw extracts from freshly harvested earthworms HE

A first batch of freshly harvested earthworms (HE) was promptly rinsed with distilled water and dried using Joseph paper. The fresh weight of the HE batch (FW_HE = 100g) was recorded before the worms were euthanized by submersion in a container containing pure methanol. The mixture was then evaporated in an oven at 60°C for three days until the weight stabilized. Following evaporation, the dried earthworms and the excretions released by the methanol treatment were ground into a powder, and the dry weight (DW_HE = 25g) was determined.

The crude extract was prepared by maceration in successive baths of distilled water, each approximately 50 mL, conducted at 60°C. Each bath was allowed to macerate for 24 hours. The brown supernatant from each maceration was gently recovered, and the process was repeated until the final macerate became transparent. The successive macerates were pooled and filtered twice—first through glass wool and then through Whatman paper. The filtrate was then evaporated in the oven at 60°C, and the dry weight of the resulting residue (DW_CEx = 11g) was determined. This dry residue, composed entirely of water-soluble substances, constitutes the crude extract from the HE batch, which will be used to prepare bacterial culture media by successive dilutions in distilled water.

#### Preparation of crude extracts from fasted earthworms FE

A second batch of earthworms (FE) is thoroughly rinsed with deionized water and then fasted for 8 to 10 days in earth pots, which are devoid of food and soil. To prevent dehydration, the worms are wrapped in moistened tissue. Every two days, the FE batch is cleaned of its casts and transferred to a new pot. This process is repeated 4 to 5 times until the casts are nearly exhausted. Additionally, stereoscopic examination is employed to confirm that the digestive tracts of the earthworms are nearly empty.

At the conclusion of the fasting period, the batch of earthworms is rinsed with distilled water and subsequently dried using blotting paper. The fresh weight of the FE batch is adjusted to match that of the HE batch, ensuring FW_FE = 100g. The crude extract from this second batch (FE) is prepared by maceration, following the same protocol used for the first batch (HE). As with the HE batch, the dry weight of the FE batch (DW_FE = 19g) and the soluble residue of its crude extract (DW_CEx = 13g) are then determined.

#### Preparation of culture media based on crude extracts of earthworms (HE and FE)

Ten grams of dry weight crude earthworm extract, from either the first batch (HE) or the second batch (FE), are dissolved in 200 mL of distilled water to prepare the HE and FE stock solutions. These stock solutions will serve as the basis for preparing the bacterial culture media through successive dilutions. Given that the pH of the HE or FE stock solution is acidic (pH = 4), it will be adjusted to a neutral pH, in alignment with the pH of conventional control media.

The 200 mL of stock solution is then divided into two 100 mL portions. The first portion (100 mL) will be used to prepare culture medium C1 at 5%. The second portion (100 mL) is adjusted to a final volume of 200 mL with distilled water, bringing it to a neutral pH. After thorough homogenization, this solution is divided again. The first 100 mL will constitute culture medium C2 at 2.5% (Fd = 2X), while the second 100 mL will be further diluted in the same manner to yield culture medium C3 at 1.25% (Fd = 4X).

### Preparation of conventional media

The effect of culture media (C1, …, C8) prepared based on earthworm extract (HE, FE) on the growth of fungi will be compared to that of the conventional medium: PDA and SAB.

- PDA (Potato dextrose agar): Potato extract 4 g/L, Glucose 20 g/L.
- SAB (Sabouraud): Peptones 10 g/L, Glucose 40 g/L.

### Evaluation of fungi growth

#### Inoculation of media and growth parameter

Mycelial growth is estimated after 4 days of incubation at 25°C by measuring the average diameter of the colonies. In each medium, three Petri dishes are used. Mycelial discs, taken from 5-day-old cultures on the PDA medium, are placed in the center of the Petri dishes containing the culture media to be tested.

#### Effect of media on sporulation

A 5 mL of physiological water was added to the surface of the fungi in solid culture media. 1 mL of the suspension was recovered in tubes. After stirring in the vortex, the spores are counted using a Malassez cell at the rate of 3 counts per suspension (Benslim et al. 2016). The number of spores/mm2 in the control medium is compared to that counted in media prepared from earthworm extracts (HE or FE).

#### Effect of Media on Spore Germination

A 0.5 mL aliquot of spore suspension at a concentration of 10^7^ spores/mL was added to a series of tubes containing 5 mL of the respective liquid media. The tubes were prepared in triplicate and incubated for 24 hours at room temperature. Following incubation, the number of germinated spores was quantified using a microscope and a Malassez counting chamber. The spore germination rate in the control medium was compared to that in media prepared from earthworm extracts (HE or FE).

### Statistical analysis

The data on fungal diameter, spore count, and spore germination were analyzed using ANOVA (SPSS version 22) based on the experimental models. Statistical significance was determined at a 0.05 probability level. To compare the means, the Tukey’s multiple comparison test was employed. All data are presented as the mean ± standard error.

## Results

The diameters of the fungi *Trichoderma* sp., *Melanocarpus* sp., *Acaulospora* sp., were measured, after 4 days of incubation in the different culture media prepared from crude extracts of earthworms HE (earthworms freshly harvested) and FE (earthworms fasted) at different concentrations (C = 5%, …, C 8 = 0.04%), and in the control culture media PDA and SAB.

### Effect of crude extract of freshly harvested earthworms (HE) and earthworms previously deprived of soil and food (FE) on mycelial growth

The results of mycelial growth in control media (PDA and SAB) and media formulated from freshly collected earthworms (HE) and earthworms deprived of soil and food (FE). The three fungi tested showed positive mycelial growth on the HE and FE culture media. However, this growth differs from one species to another and from one concentration to another. In the HE medium, the mycelial diameter of the fungus *Trichoderma* sp. increases steadily to a maximum value of 8.25 cm at the concentration C6=1.6%. Growth was significantly (p<0.05) higher in C5, C6, C7 (with diameters of 8.11, 8.25, and 8.12 cm, respectively) compared to other concentrations. In FE media, mycelial growth continued up to C8=0.04% and was significantly (p<0.05) high at C7 and C8 concentrations where the diameters were 7.98 and 8.33 cm, respectively. The diameter of the mycelium on these concentrations of HE and FE crude extracts is always close to that of PDA (8.01 cm). On the other hand, the growth on the SAB control medium (5.67 cm) was, significantly (p<0.05), very low compared to that in the HE and EF. Overall, if we compare the concentrations of extracts that gave high growth rates C6 for HE and C8 for FE, we see that the effect of the FE extract is greater than that of HE.

In the case of the fungus *Melanocarpus* sp., it was found that mycelium development stops from dilution C7=0.08% for both types of HE and FE extracts. The diameter of the mycelium increased from C1, which represents the concentrated medium containing 5% of the extract (HE or FE), to the concentration C5 for HE and C4 for FE where the growth noted was maximum about 7.90 cm. We found that in the case of this fungus, the effect of HE on mycelial growth is greater than that of EF. No significant differences were observed between the different concentrations of HE media and the control media. In the FE extract, the C4 concentration gave an average diameter of 7.95 cm which was significantly (p<0.05) high. In the PDA and SAB control media, where the diameter was 7.43 cm and 6.5 cm respectively, the amount of nutrient was 2.4% and 5%, respectively, compared to only 0.62% in the medium prepared from crude earthworm extracts.

Unlike *Trichoderma* sp. and *Melanocarpus* sp., the fungus *Acaulospora* sp. showed low growth in all concentrations of both types of HE and FE media. This low growth rate is also observed in conventional PDA and SAB control environments. However, in the FE medium, the mycelial growth in the concentration C7, which is 5.29 cm, was more or less close to that in the PDA medium whose diameter was 5.66 cm. For the HE medium, the diameter of the mycelium increases as the concentration of extract decreases, reaching a growth of 4.54 cm at the concentration C6 = 0.16%, then, it begins to decrease at low concentrations. In the FE medium, the diameter of the mycelium increases slowly from the concentration C1 to C7. Of all the concentrations of HE and FE tested, C5, C6 and C7 are the best and promoted significantly (p<0.05) the mycelium development.

### Effect of crude extract of earthworms on the dissemination and the germination of fungal spores

The results of the enumeration of spores released on solid media and spores germinated in liquid media for the three studied fungi *Trichoderma* sp., *Melanocarpus* sp., and *Acaulospora* sp., are shown in Tables 2, 3 and 4, respectively. Based on the results of the study, Table 2 shows that the number of spores released by *Trichoderma* in solid HE media reaches its highest value in the concentration C2 and then begins to decrease. In C2 and C3 media, the number of spores is significantly high, 188.33 × 10^4^ and 176.67 × 10^4^ spores mL^-1^, respectively, compared to the rest of the dilutions, and close to that of the PDA medium (186.67 × 10^4^). In the case of FE, the concentration range from C1 to C5 gives a higher average number of spores released compared to C6 to C8 concentrations and SAB medium, but significantly comparable with PDA medium. The number of spores germinated at the dilution C5 of HE extract was significantly high (49.33 × 10^2^ spores sprouted mL-1) compared to other dilutions and compared to controls PDA (38 × 10^2^) and SAB (10.33 × 10^2^). As for the FE medium, the C6 dilution gives a significantly better spore germination (53.33 × 10^2^) compared to other the dilutions and compared to PDA and SAB. Compared to conventional SAB medium, HE media, with the exception of the C1, give higher spore gemination rates.

**Table 1:**
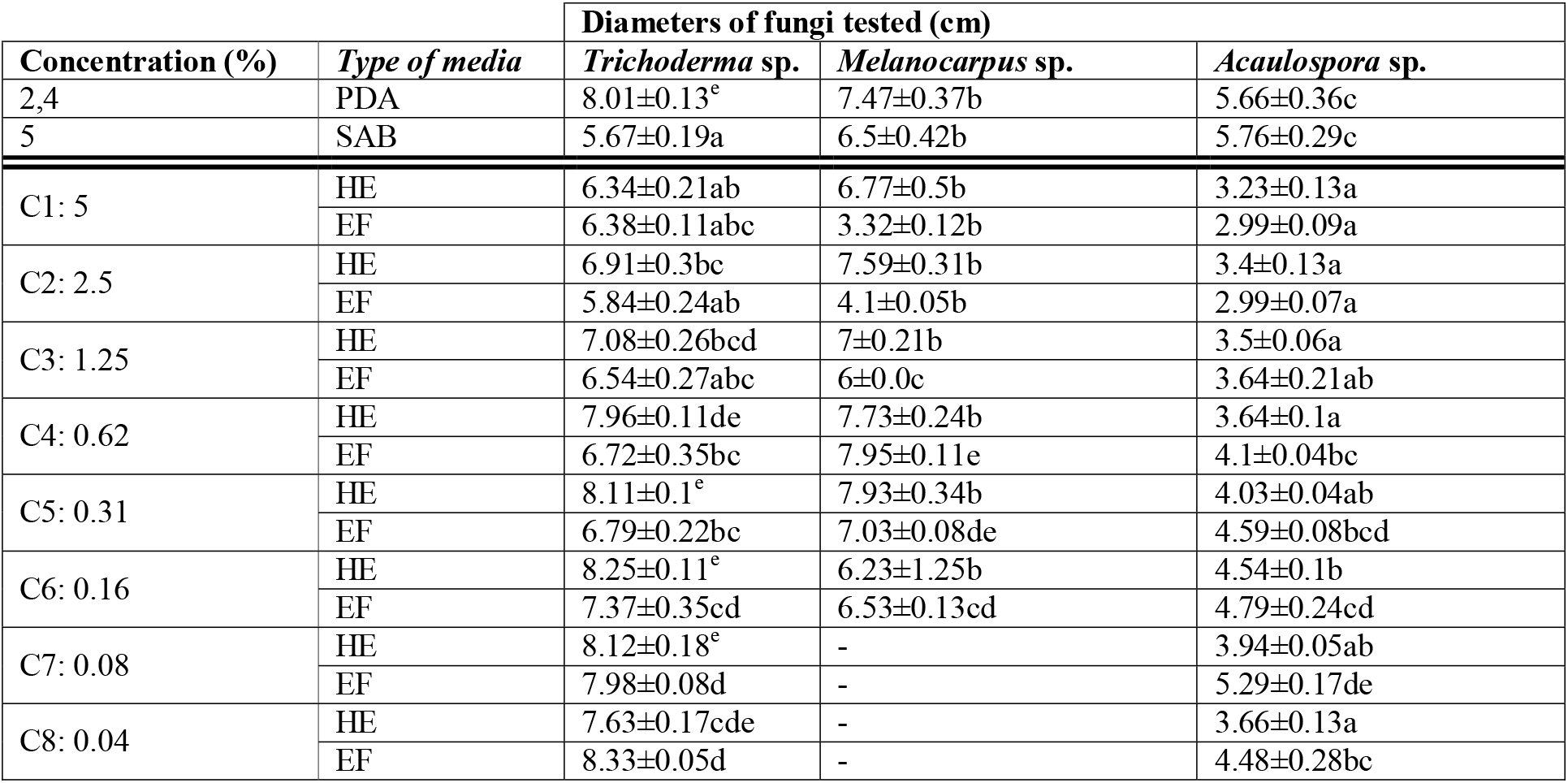
Diameter (cm) of *Trichoderma* sp., *Melanocarpus* sp., and *Acaulospora* sp. after 4 days of incubation in various culture media formulated from the crude extracts of freshly harvested earthworms (HE). and those prepared from the crude extracts of the FE earthworms (fasted earthworms) at different concentrations (C1=5%, …, C8=0.04%), and in the control culture media PDA and SAB. Growth is expressed as the mean diameter (± standard error) at each concentration. The letters indicate statistically significant differences between the culture media at a 0.05 confidence level, as determined by the Tukey’s Test (p ≤ 0.005).

| Concentration (%) | Type of media | Diameters of fungi tested (cm) |  |  |
| --- | --- | --- | --- | --- |
|  |  | *Trichoderma* sp. | *Melanocarpus* sp. | *Acaulospora* sp. |
| 2,4 | PDA | 8.01±0.13<sup>e</sup> | 7.47±0.37b | 5.66±0.36c |
| 5 | SAB | 5.67±0.19a | 6.5±0.42b | 5.76±0.29c |
| C1: 5 | HE | 6.34±0.21ab | 6.77±0.5b | 3.23±0.13a |
| | EF | 6.38±0.11abc | 3.32±0.12b | 2.99±0.09a |
| C2: 2.5 | HE | 6.91±0.3bc | 7.59±0.31b | 3.4±0.13a |
| | EF | 5.84±0.24ab | 4.1±0.05b | 2.99±0.07a |
| C3: 1.25 | HE | 7.08±0.26bcd | 7±0.21b | 3.5±0.06a |
| | EF | 6.54±0.27abc | 6±0.0c | 3.64±0.21ab |
| C4: 0.62 | HE | 7.96±0.11de | 7.73±0.24b | 3.64±0.1a |
| | EF | 6.72±0.35bc | 7.95±0.11e | 4.1±0.04bc |
| C5: 0.31 | HE | 8.11±0.1<sup>e</sup> | 7.93±0.34b | 4.03±0.04ab |
| | EF | 6.79±0.22bc | 7.03±0.08de | 4.59±0.08bcd |
| C6: 0.16 | HE | 8.25±0.11<sup>e</sup> | 6.23±1.25b | 4.54±0.1b |
| | EF | 7.37±0.35cd | 6.53±0.13cd | 4.79±0.24cd |
| C7: 0.08 | HE | 8.12±0.18<sup>e</sup> | - | 3.94±0.05ab |
| | EF | 7.98±0.08d | - | 5.29±0.17de |
| C8: 0.04 | HE | 7.63±0.17cde | - | 3.66±0.13a |
| | EF | 8.33±0.05d | - | 4.48±0.28bc |

**Table 2:** The number of spores released and germinated (± standard error) by the fungus *Trichoderma* sp. respectively after incubation for 4 days in solid media and 24 hours in liquid media. Culture media prepared with varying concentrations of earthworm (*A. molleri*) extracts, ranging from C1 (5%) to C8 (0.04%), were derived from both freshly harvested earthworms (HE) and earthworms previously deprived of food and soil (FE). Conventional control media: PDA (2.4%) and SAB (5%). The letters indicate statistically significant differences between culture media at a 0.05 confidence level, as determined by the Tukey test (p ≤ 0.005).

|  |  | *Trichoderma* sp. |  |
| --- | --- | --- | --- |
| Concentration (%) | Type of media | Number of released spores $10^4 \text{ mL}^{-1}$ | Number of germinated spores $10^2 \text{ mL}^{-1}$ |
| 2,4 | PDA | 186.67±10.93b | 38±9.02 ABC |
| 5 | SAB | 65±5.77a | 10.33±2.33 ab |
| C1: 5 | HE | 106.67±29.2a | 7.33±1.76 a |
| | FE | 165±7.64b | 18.67±5.81 ab |
| C2: 2.5 | HE | 188.33±4.4b | 22.33±1.2 abc |
| | FE | 168.33±10.14b | 26.67±2.67abc |
| C3: 1.25 | HE | 176.67±10.93b | 30±7.5 abc |
| | FE | 158.33±13.64b | 32±8.33 abc |
| C4: 0.62 | HE | 101.67±13.64a | 41.33±3.53abc |
| | FE | 176.67±7.26b | 42.67±7.42abc |
| C5: 0.31 | HE | 100±5.77a | 49.33±1.33 bc |
| | FE | 153.33±6.01b | 46.67±7.42 bc |
| C6: 0.16 | HE | 100±13.23a | 44±10.58 b |
| | FE | 78.33±1.67a | 53.33±2.67 c |
| C7: 0.08 | HE | 60±7.64a | 30.67±13.13 abc |
| | FE | 91.67±16.91a | 49.33±5.81 bc |
| C8: 0.04 | HE | 75±10.41a | 30.67±7.05abc |
| | FE | 51.67±7.26a | 38.67±5.81abc |

**Table 3:** The number of spores released and germinated (± standard error) by the fungus *Melanocarpus* sp. respectively after incubation for 4 days in solid media and 24 hours in liquid media. Culture media prepared at varying concentrations, ranging from C1 (5%) to C8 (0.04%), were formulated using extracts from freshly harvested earthworms (*A. molleri*) (HE) and from earthworms previously deprived of food and soil (FE). Conventional control media: PDA (2.4%) and SAB (5%). The letters indicate statistically significant differences between culture media at a 0.05 confidence level, as determined by the Tukey test (p ≤ 0.005).

|  |  | *Melanocarpus* sp. |  |
| --- | --- | --- | --- |
| Concentration (%) | Type of media | Number of released spores $10^4 \text{ mL}^{-1}$ | Number of germinated spores $10^2 \text{ mL}^{-1}$ |
| 2,4 | PDA | 230±20.82b | 29.33±5.81bc |
| 5 | SAB | 121.67±10.93a | 14.67±4.81ab |
| C1: 5 | HE | 143.33±19.22b | 13.33±3.53 ab |
| | FE | 140±18.93b | 18.67±7.05abc |
| C2: 2.5 | HE | 198.33±51.99b | 10.67±2.67 ab |
| | FE | 213.33±34.68b | 24±8.33 abcd |
| C3: 1.25 | HE | 225±28.87b | 18.67±4.81 ab |
| | FE | 208.33±29.49b | 24±6.93 abcd |
| C4: 0.62 | HE | 155±25.98b | 33.33±4.81 bc |
| | FE | 175±30.55b | 46.67±4.81 d |
| C5: 0.31 | HE | 143.33±17.64b | 54.67±10.41 c |
| | FE | 155±22.91b | 41.33±4.81cd |
| C6: 0.16 | HE | 133.33±6.01b | 21.33±7.05 ab |
| | FE | 143.33±18.78b | 33.33±2.67bcd |
| C7: 0.08 | HE | - | - |
|  | FE | - | - |
| C8: 0.04 | HE | - | - |
|  | FE | - | - |

**Table 4:**
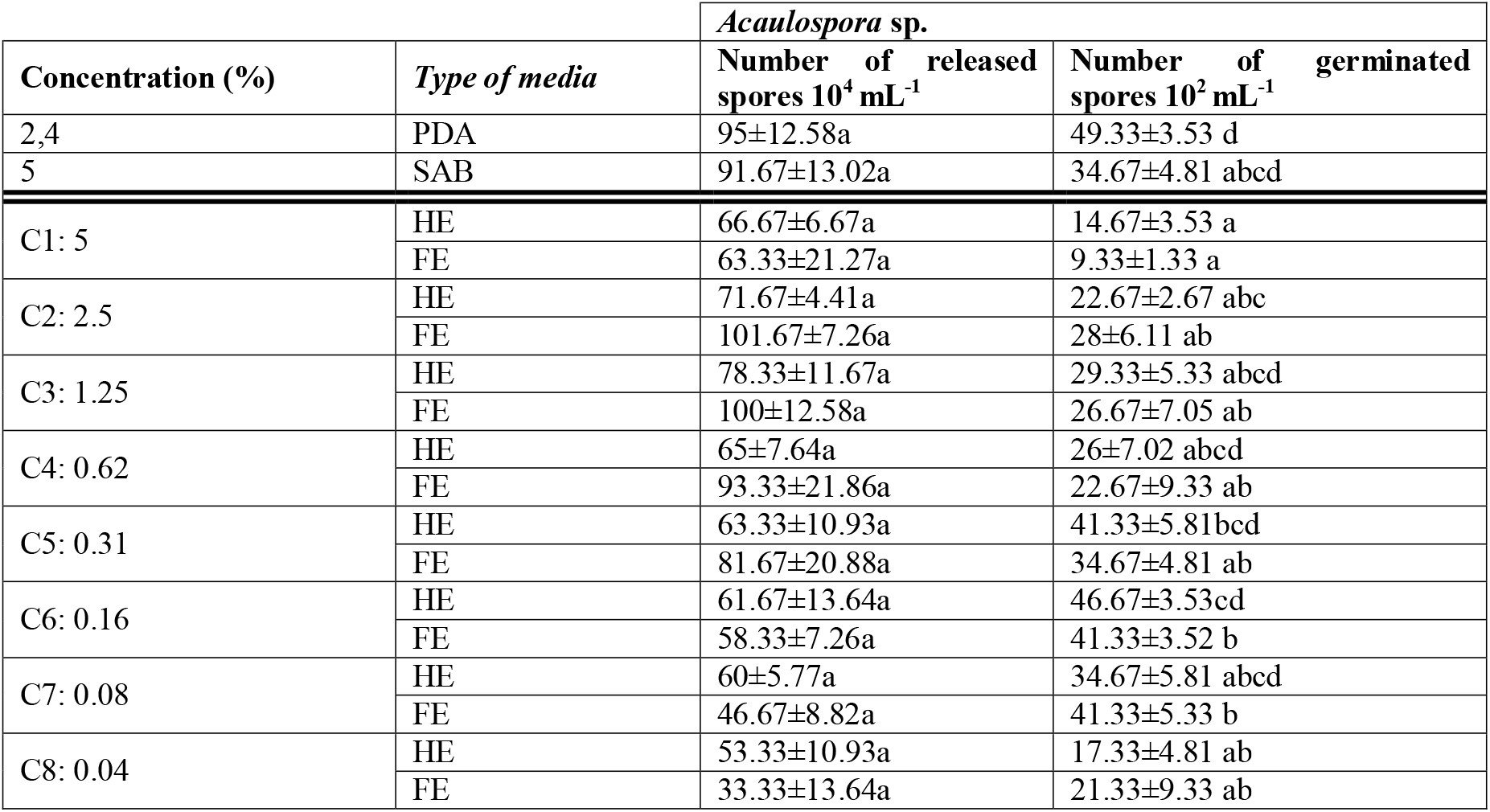
The number of spores released and germinated (± standard error) by the fungus *Acaulospora* sp. respectively after incubation for 4 days in solid media and 24 hours in liquid media. Culture media with varying concentrations (C1: 5% to C8: 0.04%) of extracts from *A. molleri* earthworms, including those freshly harvested from the field (HE) and those previously deprived of soil and food (FE). Conventional control media: PDA (2.4%) and SAB (5%). The letters indicate statistically significant differences between culture media at a 0.05 confidence level, as determined by the Tukey test (p ≤ 0.005).

In the fungus *Melanocarpus* sp., Table 3 shows that, with the exception of C7 and C8, all concentrations prepared based on HE or FE extracts, allowed the production of a significantly high number of spores compared to the control medium SAB. However, the number of spores counted in these media is comparable to that of the control medium PDA. The number of *Melanocarpus* sp. spores was significantly elevated in C5 medium (54.67 × 10 ^2^ germinated spores mL-1), followed by C4 (33.33 × 10^2^). In the rest of the concentrations, the difference was not significant. Statistically, germination in HE media is superior or similar to the control medium SAB. As in HE media, the number of spores germinated in the FE extract is always higher than that in the SAB medium (14.67 × 10^2^). The C4 concentration showed the highest (46.67 × 10 2), followed by C5 (41.33 × 10 2) and the rest of the dilutions. The control medium PDA gives a germination rate higher than that of the SAB medium and significantly comparable to that of the HE and FE media at different concentrations.

In *Acaulospora* sp. (Table 4), no significant differences in spores released were observed between media prepared from crude earthworm extracts (HE or FE) and control media SAB and PDA. The number of spores germinated (46.67 × 10^2^) in the C6 concentration of the HE extract was significantly higher compared to the other concentrations and the SAB medium (34.67 × 10^2^), but close to that recorded in the PDA (49.33 × 10^2^). In the case of FE, both C6 and C7 resulted in a higher number of spores (41.33 × 10^2^) than the number noted in the other dilutions and in the SAB medium, but significantly similar to the PDA medium. For both types of HE and FE extracts, the SAB medium gives a germination close to the media at the concentration range C2-C5 and C8 and lower compared to the concentration C1. Overall, it is interesting to note that, the behaviors of the 3 fungal strains in the two extracts HE and FE at different concentrations are similar with minor differences. Therefore, the intestinal contents of earthworms have no noticeable effect. The nutrients and growth factors of the extracts would be mainly intrinsic to the various tissue structures of the earthworms. The concentrations that allowed maximum growth of fungi are between C4=0.6% and C8=0.04% of HE and FE extracts. This amount is 4 to 100 times lower than the usual amount of nutrients from conventional control PDA and SAB media, respectively 2.4% and 5%. The evolution of mycelial growth as a function of HE and FE concentrations is almost parallel to that of the germinated spores. Generally, it is found that there is a positive correlation between spore germination rates and mycelial growth. On the other hand, the evolution of released spores is quite different from that of the evolution of germinated spores. Sporulation is favored at higher concentrations between C1=5% to C4=0.62% regardless of the type of extract.

## Discussion

In conclusion, this study demonstrates that crude extracts from earthworms, both freshly harvested from the field (HE) and those deprived of soil and food for 10 days (FE), significantly promote fungal growth at substantially lower nutrient concentrations compared to conventional media, which typically contain 2.4% and 5% nutrient concentrations. These findings suggest that media formulated with HE and FE earthworm extracts offer a more diverse array of nutrients and growth-promoting substances, thus presenting promising alternatives for enhancing fungal cultivation.

The results of this study lead to the conclusion that the contents of the earthworm digestive tract play a negligible role in enhancing fungal growth. If the digestive contents were a significant factor, one would expect the extract from freshly harvested (HE) earthworms to exhibit greater efficacy than the extract from earthworms that had been deprived of soil and food for 10 days (FE), which have an empty digestive tract. Prior assessments have shown that the digestive tract content represents approximately 6 g per 25 g of dry worm weight (24%) (Yakkou et al., 2021b), predominantly composed of soil detritus mixed with organic matter. This substantial amount, however, is unlikely to substantially contribute to the nutrient enrichment of the HE extract. Therefore, we affirm that the primary source of nutrients and growth factors is likely endogenous to the tissue structures of the earthworms themselves. Consistent with the findings of our previous work, the contents of the digestive tract did not appear to influence microbial growth, as the efficacy of the FE extract (with an empty digestive tract) was comparable to that of the HE extract (with a full digestive tract) (Yakkou et al., 2021a, b).

Our previous study demonstrated that crude extracts from freshly harvested earthworms, those deprived of soil and food for 10 days, and cutaneous excreta, all significantly promote bacterial growth (Yakkou et al., 2021a, 2021b). Collectively, these findings indicate that the decomposition of earthworm bodies, alongside the continuous production of cutaneous excreta, enriches the soil with high-value nutrients and growth factors. These additions play a crucial role in supporting the activity and development of plant growth-promoting rhizobacteria (PGPR) and PGPF, both of which are vital for enhancing soil fertility and promoting plant growth.

The protein content in earthworms varies significantly across species, ranging from 32.6% to 67.2% (Damayanti et al. 2008; Julendra 2003), and includes ten essential amino acids and ten non-essential ones. Isoleucine is the most prevalent essential amino acid, comprising 41.98% of the dry weight, while glutamic acid is the dominant non-essential amino acid at 1.52% of dry weight (Istiqomah et al. 2009; Jeyanthi et al. 2016). Earthworm dry matter additionally contains 7–10% fats, 8–20% carbohydrates, and 2–3% minerals (Ghatnekar et al. 1995).

The fungi *Trichoderma* sp., *Melanocarpus* sp., and *Acaulospora* sp. are recognized for their roles in soil fertility enhancement and plant growth promotion, both directly and indirectly. These fungi significantly inhibit the growth of phytopathogens and modulate plant growth rates. Known as plant growth-promoting fungi (PGPF), they produce extensive networks of mycelial filaments that explore large soil volumes, enhancing nutrient recovery beyond the capacity of plant root hairs (Clark 1997; Naseby et al. 2000; Siddiqui and Shaukat 2004; Zin and Badaluddin 2020). Their crucial role, particularly in phosphate solubilization, enhances phosphorus bioavailability in the rhizosphere (Souchie et al. 2006). Furthermore, depending on environmental context, these fungi contribute to the assimilation of additional nutrients such as nitrogen, potassium, calcium, and trace elements (Imtiaz et al. 2016). Given their ability to colonize deep soil layers, fungal mycelia also assist in plant water uptake and resistance to drought stress (Bano and Ashfaq 2013).

Our findings that earthworm-derived media stimulate fungal spore formation are noteworthy, as spores represent the primary means for fungal dispersal, facilitating the colonization of new ecological niches and ensuring reproductive success under adverse conditions (Dahlberg and Etten 1982). Sporulation, a complex differentiation process involving somatic-to-reproductive cell transformation, typically initiates when hyphal growth decelerates (Dahlberg 1982), aligning with our observed results. Concentrated earthworm extract media promoted spore production, while more dilute media favored spore germination and mycelial expansion.

Fungal spore germination, marked by the emergence of a germ tube, is contingent on nutrient availability—primarily amino acids, sugars, and nucleosides. Similar to hyphal elongation, vesicles accumulate at the plasma membrane to supply structural materials to the growing wall, forming a specialized structure known as the Apical Vesicle Complex at the germ tube tip (Kye 2008). The primary mycelium, branching from the germ tube, grows into a hyphal network, resulting in a strong positive correlation between spore germination and hyphal development under similar culture conditions (Guillaume 2018).

Efforts to establish economical culture media alternatives to PDA for fungal cultivation have so far yielded growth outcomes that are either comparable or inferior to PDA’s (Jadhav et al. 2018). Notably, in our study, the concentration of crude earthworm extract necessary for optimal fungal growth was 4–100 times lower than nutrient levels in standard PDA (2.4%) and SAB (5%) media, marking a promising advance. Despite extensive research aimed at developing alternative media for soil microorganism isolation and cultivation, results to date have been limited, often due to the inadequacy of the media in meeting microbial requirements (Vartoukian et al. 2010; Uthayasooriyan et al. 2016; Jadhav et al. 2018). The promising outcomes of this study thus motivate further research within our laboratory to develop a novel earthworm-extract-based culture medium, potentially surpassing conventional media in efficacy for isolating and culturing soil bacteria and fungi.

## Acknowledgements

This study was conducted with the invaluable support of the National Center for Scientific and Technical Research (CNRST) through the Research Excellence Scholarships Program.

We extend our sincere gratitude to Mr. Jorge Dominguez for his expertise and assistance in conducting the molecular identification of the earthworm species.

## Declarations

### Compliance with Ethical Standards

The authors affirm that all ethical standards were strictly adhered to in the conduct of this research, and that no human participants or animals were involved.

### Consent for Publication

All authors have provided their consent for publication of this work.

